# *SurfacOmics:* an R shiny application integrating variable Feature Selection for gene biomarker discovery using Elastic-Net Regularization

**DOI:** 10.1101/2025.06.12.659259

**Authors:** Sadhana Tripathi, Priya Kajla, Bilal Ahmed Abbasi, Kota Priya, Aubrey Bailey, Binuja Varma

## Abstract

Affordable sequencing technologies have resulted in a rapid rise in genomic and proteomic data. As a result, a massive amount of data is being analyzed, and the outcomes must be summarized as relevant clinical biomarkers. One of the major challenges is enabling wet-lab researchers to make meaningful inferences from the data, even in the absence of expertise in pipeline development and statistical training. We present a user-friendly R shiny application, *SurfacOmics*, which allows the user to perform biomarker identification via penalized regression algorithms. It also enables researchers to choose a crucial binary variable, such as treatment or group, from the study design metadata to anticipate potential biomarkers. We have introduced a novel concept of scoring scheme to characterize potential biomarkers for prediction, prioritization, and ranking. The tool integrates Gene Ontology information for sub-cellular localization together with a manually curated knowledgebase to provide valuable insights on biological processes, molecular functions and antibody resources, offering a comprehensive view in a single interface.

## Introduction

Over the past decade, large-scale omics data analysis has attracted significant attention for its potential to elucidate complex biological phenomena, disease markers, and precision medicine (1). The discovery of biomarkers via genomics, proteomics, and metagenomics data remains a major goal in translational medicine and functional genomics (2,3). Identification of new biomarkers would allow clinicians and researchers to meaningfully interpret underlying disease mechanisms and could often aid in early definitive diagnosis, prognosis, and risk prediction of diseases (4). As an increasing amount of complex data is generated, the integration of sophisticated algorithms, pipeline development, and thorough statistical knowledge become the true limiting factors in identifying pertinent biomarkers (5,6).

Machine learning techniques have been broadly used to investigate disease mechanisms, discover predictive biomarkers, and develop stable risk prediction models on multi-omics data (7). Glmnet-based elastic net regression algorithms offer efficient modeling of high-dimensional omics data, balancing feature selection and regularization—even in scenarios with limited samples (n << p) (8,9). It is also well known for its ability to capture complex dependency patterns between the outcome and variates.

Identifying relevant biomarkers from extensive lists of genes or proteins necessitates the input of experts in different fields. This exchange of information between the data experts and the laboratory researchers is sensitive, and at times vital findings can be lost or misinterpreted during brief discussions (10). Several platforms support interactive analyses, enabling data interpretation without programming (11–14). Still, there are limited open-source tools specifically for genetic biomarker detection that are accessible without hardcore programming or command-line skills. This presents a unique opportunity to make and enhance the wet-lab researchers’ experience for discovering gene biomarkers.

We have developed *SurfacOmics*, an R Shiny application, to equip biomedical researchers with minimal programming expertise. This tool employs elastic net regression to identify biomarkers and ranks them using a manually curated knowledgebase. To help users understand the role of these biomarkers, SurfacOmics provides insights into their biological functions and pathways through Gene Ontology (GO) data. Additionally, the application includes a list of antibody resources for each biomarker to support experimental validation. We have also added a scoring system to predict, prioritize, and rank biomarkers. Lastly, the *SurfacOmics* generates detailed reports with interactive charts and tables to make evaluating biomarkers simple and efficient.

We envision *SurfacOmics* as a valuable bridge between data-driven analysis and experimental research, streamlining biomarker discovery for applications in diagnostics, therapeutics, and precision medicine.

## Methodology

The Shiny application *‘SurfacOmics*’ is implemented using R version 4.1.2 (2021-11-01) and provides an interactive interface to facilitate each data analysis step (**Figure 1)**. It performs three major operations: prediction, prioritization, and ranking, leading to identification of promising candidate biomarkers.

**Figure 1.**
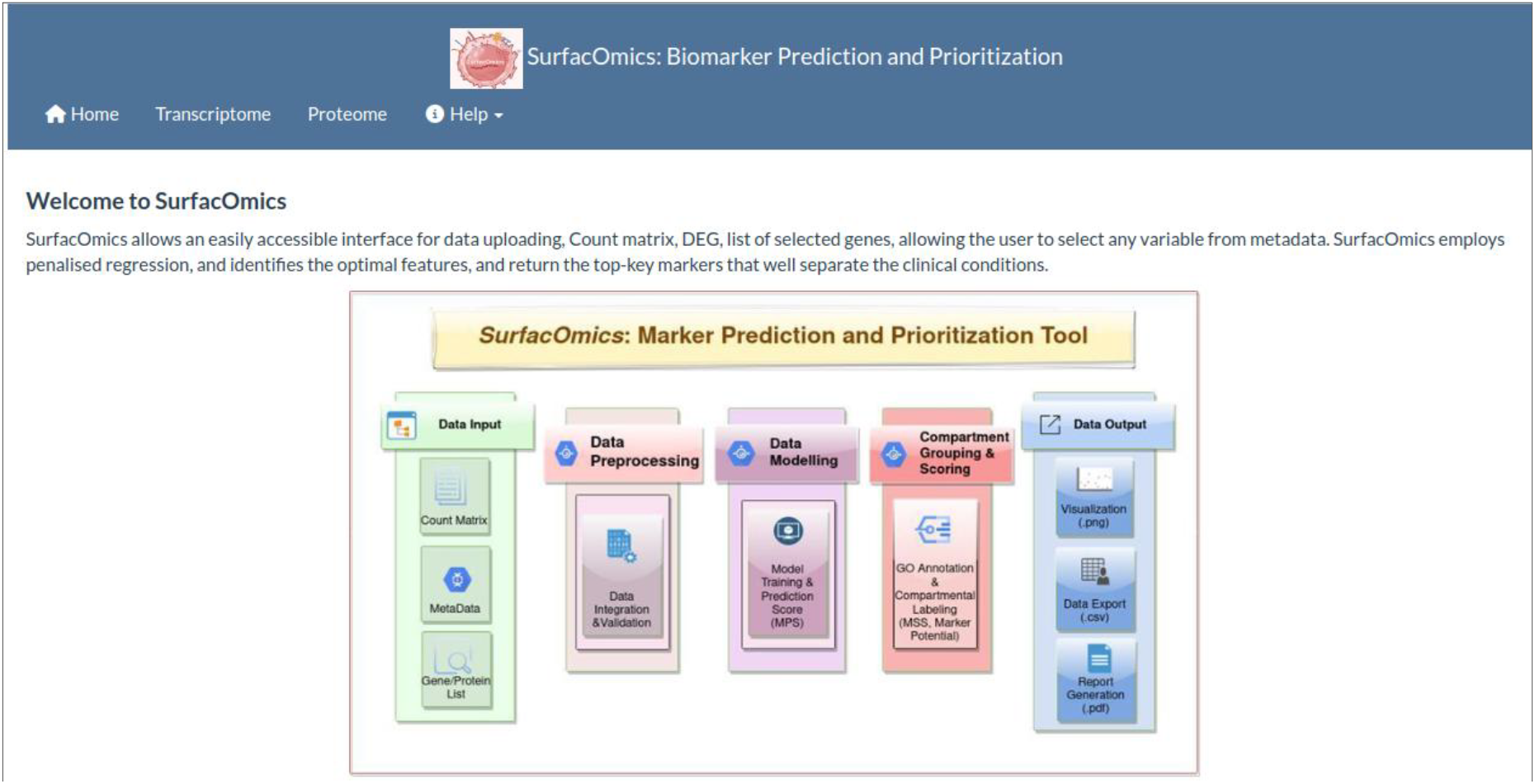
Schematic Representation of SurfacOmics’s Graphical Interface. This figure illustrates the GUI of the tool, highlighting the key components and functionalities. It provides a visual overview of the user interaction design, including essential elements such as data input, data processing, data modelling, data output, scoring as well as the operational flow.

It requires three input files from the user: a count matrix (CSV), a metadata file (CSV), and a list of selected genes or proteins (TXT). Users are provided with the option to customize and accommodate learning variables, as well as a choice to set regularization and penalization parameters for model training. Additionally, it also conducts basic sanity checks to validate the input files before proceeding with prediction operations.

### Marker Prediction

To identify pertinent biomarker candidates, *SurfacOmics* uses a general linear model (GLM) with elastic net penalization from the R package “glmnet” (15). This algorithm tries to fit a generalized linear model implementing the penalized maximum likelihood in a cyclical coordinate descent method, making it fast, accurate, and successful in optimizing the objective function for parameters (16). It uses a combination of two different penalty functions with tuning parameters which help shrink the beta-coefficients in GLM. It aims to fit linear, logistic, and multinomial Poisson and Cox regression models. The additional R package “caret” was used to fit the elastic net GLM model with a custom option for the GLMnet algorithm to identify the optimum features for tuning parameters (17).

The scarcity of experimental data poses a challenge in model training, especially when there are only two samples in each study group. *SurfacOmics* addresses this using Lasso, a machine learning technique to optimize analysis and mitigate the impact of limited sample size.

### Data Modeling

GLMnet is a fast estimation procedure combining generalized linear models with convex penalties, such as *ℓ*1 (lasso) and *ℓ*2 (ridge regression), a combination of which comprises elastic net penalties (16). Individually, ridge regression is mainly used for handling many explanatory variables and predictors. The model handles this by shrinking the corresponding coefficients of collated variables to zero, resulting in regression coefficients non-zero (18). Additionally, lasso only selects a key correlated variable and ignores the remaining variables.

The user can customize and tune the options, or hyperparameters, alpha (α) and lambda (λ). Generally, α lies 0<α<1, offering a balance between feature selection and coefficient shrinkage. This provides the contribution weights of *ℓ*1 and *ℓ*2 penalties.

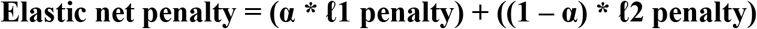

The second tuning hyperparameter, λ, controls the amount of regularization and overall weighing of the sum and strength of both penalties. The larger the λ, the coefficient shrinks toward zero. If λ is disabled or zero, the general linearized model is implemented (15).

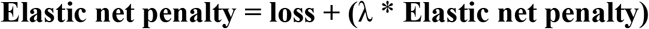

User can either choose to use the “default” or the “customize” option for α and λ parameters. Enabling the “default” option provides best-fit values determined automatically by the model on top of the user provided dataset. However, a sliding option in GUI is also provided to tweak these parameters for customization.

### Marker Prioritization

To aid biomarker prioritization, we manually curated a comprehensive knowledgebase, called SurfacTag. Sourced from Gene Ontology (GO) and UniProtKB (19), it categorizes genes and proteins based on their cellular location, with six main labels: Inside cell, Outside cell, Function Dependent, On cell surface, Cell Periphery, and Outside on cell surface. The SurfacTag knowledgebase is designed to support multiple model organisms, specifically *Homo sapiens, Mus musculus, Sus scrofa, Drosophila melanogaster, Macaca mulatta*, and *Danio rerio*. This broad support makes this tool valuable for researchers working across a variety of organisms.

One of the key challenges encountered during the creation of SurfacTag was the issue of a single gene or protein having multiple possible labels based on its different localizations under varying conditions (**Supplementary Table 1)**. To resolve this, different use cases were introduced. These use cases ensure that the database accurately assigns the predominant label or SurfacTag, providing more precise and relevant information.

**Table 1.** Potential Biomarkers identified by the *SurfacOmics* application.

| Marker Gene | Label | Marker Prediction Score | Marker Surface Score | Marker Potential | Marker Strength |
| --- | --- | --- | --- | --- | --- |
| ADAMTS17 | Cell Periphery | 100 | 4 | 1 | Strong |
| BMP8B | No information | 99 | 0 | 0.59 | Moderate |
| CBR4 | Inside cell | 30 | 1 | 0.28 | Poor |
| ENSSSCG00000045564 | No information | 6 | 0 | 0.04 | Poor |
| GPT2 | No information | 98 | 0 | 0.59 | Moderate |
| SDS | Inside cell | 16 | 1 | 0.20 | Poor |

### Ranking Biomarkers

To rank the probable biomarkers, scores have been assigned to each gene/protein calculated using predicted probability from the penalized regression model as well as prioritized labels from the annotated knowledgebase. We also calculated a cumulative score to rank predicted and prioritized markers. Additionally, for better visual representation of these scoring matrices, we have generated plots using R packages “ggplot2” and “plotly” (20). An in-app feature has been provided to export the resulting matrix and high-resolution plots in .CSV and .PNG formats, respectively.

### Marker Prediction Score (MPS)

MPS is a metric designed to rank putative markers (gene/protein) based on ML models (Lasso, GLMnet, and PLS). MPS computes the likelihood of the discriminative and distinctive features of biomarkers among sample groups (e.g., experimental/diseased conditions and groups). A high MPS score means that the identified markers differentiate well between the two groups or categories under the learning variable selected by user such as treatment, group, or condition. In addition to this, a box plot is generated to provide informed data interpretation for identified markers.

### Marker SurfacTag Score (MSS)

MSS is another key prioritizing measure employed to score genes/proteins based on their cellular localization, ease of accessibility, and assayability. Biomarkers identified on the cell surface orienting toward extracellular membranes receive higher priority, thus a higher MSS score, while biomarkers identified inside cells are assigned a low score. Based on the SurfacTag knowledgebase, MSS score is assigned to each SurfacTag (**Supplementary Table 2**). To assist in flexible decision making, a bubble plot is generated against the prioritized biomarkers.

### Marker Potential

To further improve the likelihood of optimum biomarkers, we developed “Marker Potential” as a weighted sum of MPS and MSS scores for use as a final metric in ranking identified biomarkers (**Figure 2)**. To calculate Marker Potential, MPS and MSS have been assigned distinct weights.

**Figure 2.**
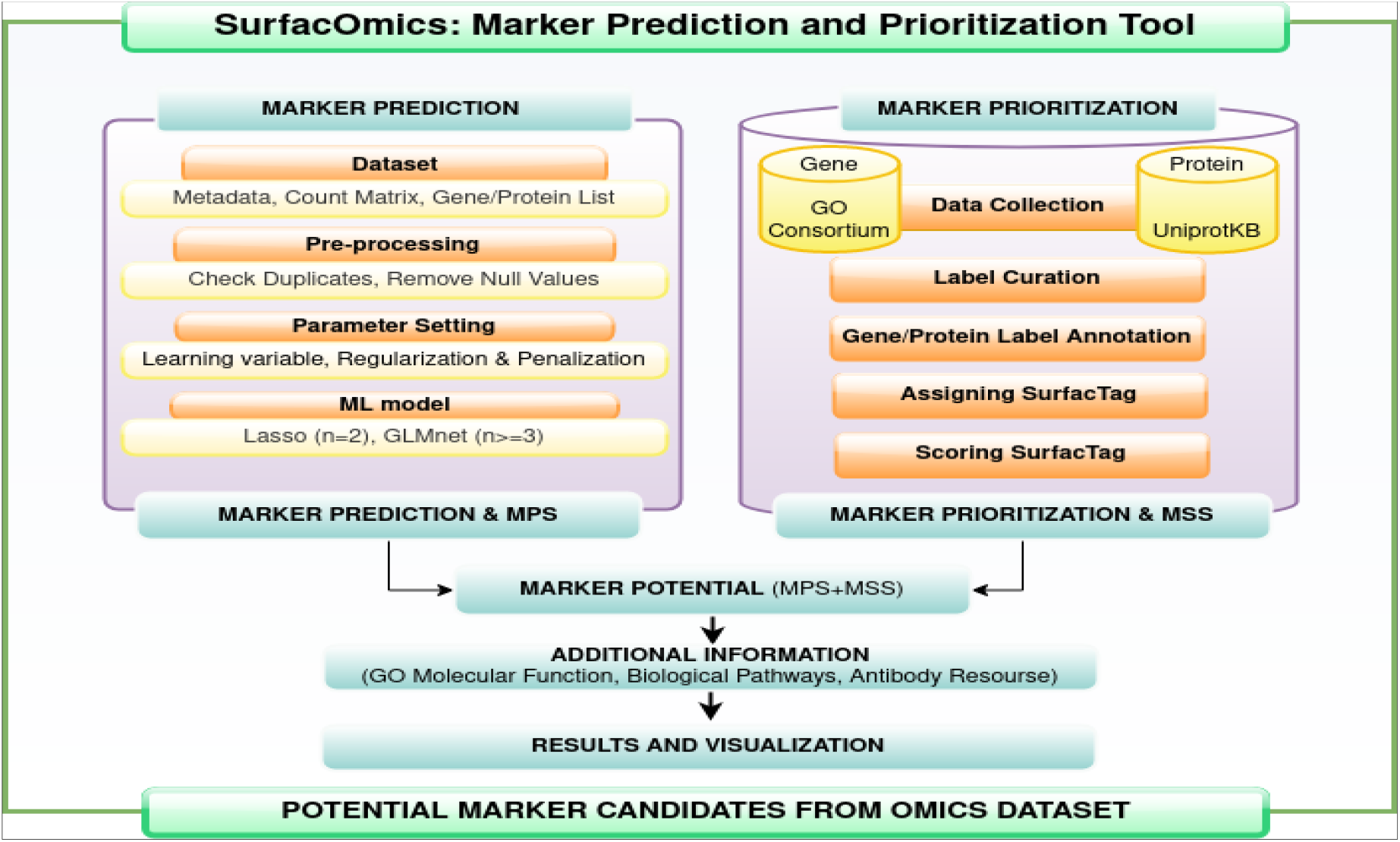
Methodology for Prioritizing Potential Biomarkers. This flowchart depicts the methodology for identifying potential biomarkers from omics datasets. It includes processes such as marker prediction, marker prioritization, scoring biomarkers, and enhancing biological relevance through the utilization of curated knowledgebases. Each step is designed to systematically refine and prioritize biomarkers for further investigation.

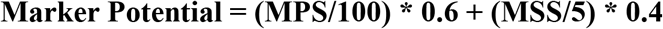

The MPS score is given a greater heuristic weighting, as all biomarker applications require classifiability while only some require those markers to be present on the cell surface. Marker Potential value ranges from 0-1. Genes and proteins with high MPS and MSS scores are assigned with relatively high Marker Potential. Similarly, genes and proteins with low MPS and MSS will yield a low Marker Potential. The Marker Potential has been assigned a magnitude and direction, ranging among weak, moderate, and strong attributes. These attributes are plotted via dot and bubble plots.

## Dataset

To demonstrate *SurfacOmics* functionalities, we use a publicly available multi-omics dataset. We shortlisted and analyzed the MIDY pig study, which focuses on diabetes mellitus in pigs (*Sus Scrofa)* with impaired insulin (*INS*) gene. This study incorporates 2-year-old female MIDY pigs (n = 4) and WT littermates (n = 5) for various omics analyses, ultimately establishing a Munich MIDY pig biobank. Multi-omics analyses were performed by comparing WT and MIDY groups (21).

## Results and Discussion

Identifying biomarkers from high-throughput omics data and performing downstream analysis like functional enrichment analysis, can be tiresome and often requires a bioinformatician or a computationally biologist (22). To enhance this process, we introduced the concept of ML based prediction, alongside SurfacTag knowledgebase-led prioritization and ranking. This combined approach enables the identification of statistically significant biomarkers using machine learning, while also prioritizing them based on cellular localization and accessibility. For a clearer interpretation, the prioritized biomarkers are ranked using a weighted score that integrates both the prediction score (MPS) and the SurfacTag score (MSS), resulting in a single value referred to as “Marker Potential”. Additionally, we have included graphical representations and plots to simplify the interpretation of the results.

This R-shiny based user-friendly application hosted at 0.0.0.0, allows users to explore the dynamics of omics dataset. Users have the flexibility to select any binary learning variable from the metadata to analyse various conditions in their study design. Users can customize regularization parameters according to their preferences or opt for default settings, where the model automatically tunes the regularization parameters based on the dataset.

Additionally, to provide a comprehensive view, this application includes detailed information on the molecular functions and biological processes associated with each gene and protein. It also offers procurement details for suitable antibodies to facilitate experimental validation, including direct hyperlinks or kit and reagent information from major suppliers (https://www.antibodyresource.com). *SurfacOmics* supports not only *Homo sapiens* (Human) but five other model organisms-*Macaca mulatta* (Rhesus monkey), *Mus musculus* (Mouse), *Sus scrofa* (Pig), *Danio rerio* (Zebrafish), *Drosophila melanogaster* (Fruit fly). Researchers can utilize data from either proteomic or transcriptomic experiments within any of these model organisms to identify, predict, prioritize, and annotate potential marker genes and proteins. Additionally, a comprehensive report, along with detailed analysis, is provided to aid in the interpretation of the results.

## Case Study

We demonstrate the capability of *SurfacOmics* application, with the help of a publicly available MIDY-pig dataset by identifying, prioritizing, and ranking markers from their transcriptomics and proteomics data.

Upon dataset submission and parameters setting, the application performs data sanity checks. The model is then trained using a selected learning variable as well as regularization and penalization parameters. As demonstrated in **Figure 3**, the learning variable is set to “Genotype” whereas regularization and penalization parameters are set to “Custom”, alpha=0.8 and lambda=0.1, respectively. Following model training and annotation, the application presents three distinct scoring matrices. Each matrix is accompanied by a scoring scheme and respective plots. These matrices showcase prediction and prioritization scores for each identified biomarker, ranked based on their significance as potential biomarkers. These scoring results (MPS, MSS and Marker potential) are summarized in a tabular format as shown in **Table 1**.

**Figure 3.**
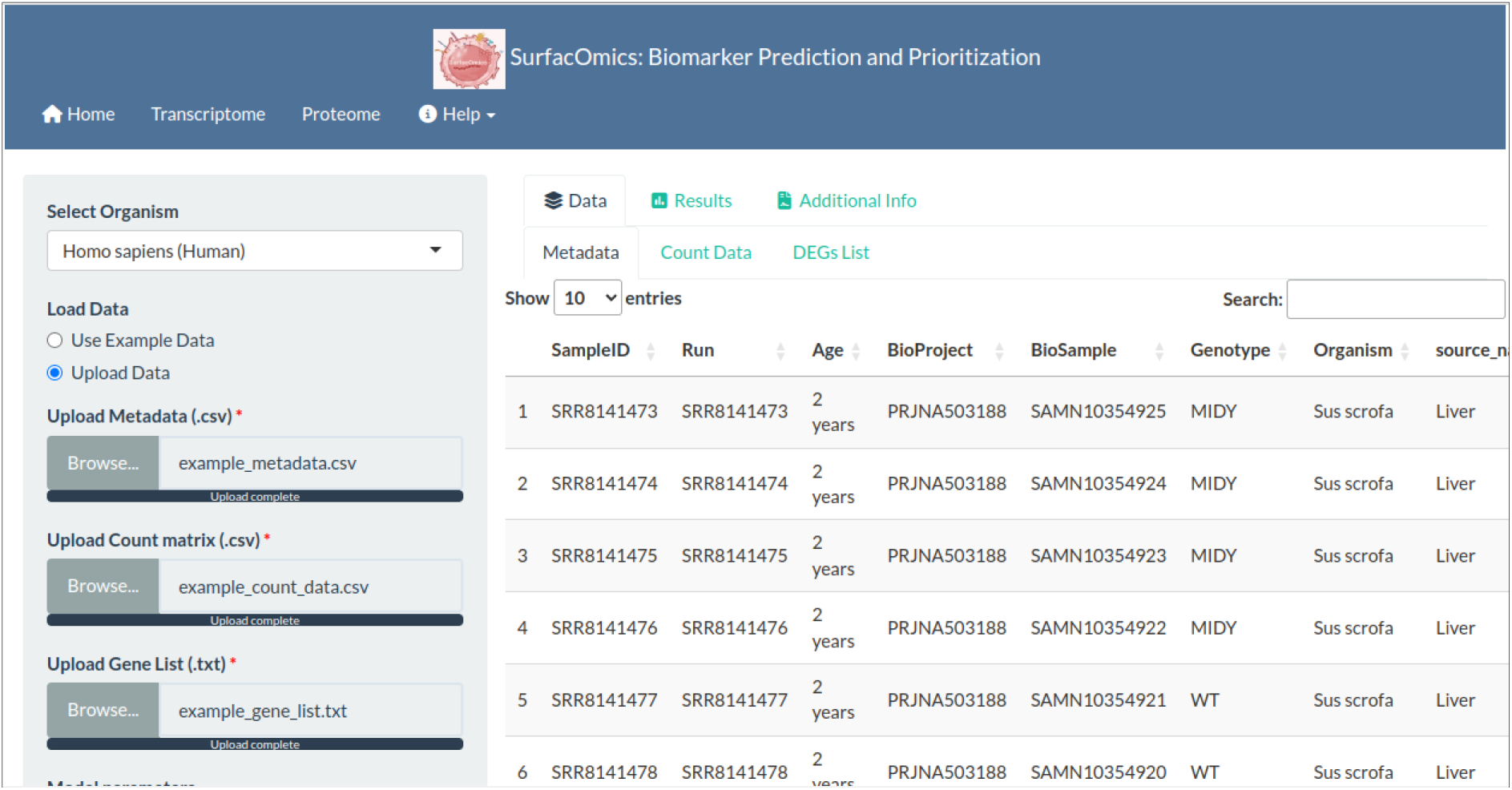
Example Display of *SurfacOmics* Interface with Sample Data. This figure presents an example display of the tool’s GUI, illustrating its user-friendly interface designed for gene biomarker discovery. The application integrates metadata, count matrix, and gene expression analysis, allowing users to efficiently explore and identify potential biomarkers. The interface provides various options to customize and streamline the analysis process.

**Figure 4.**
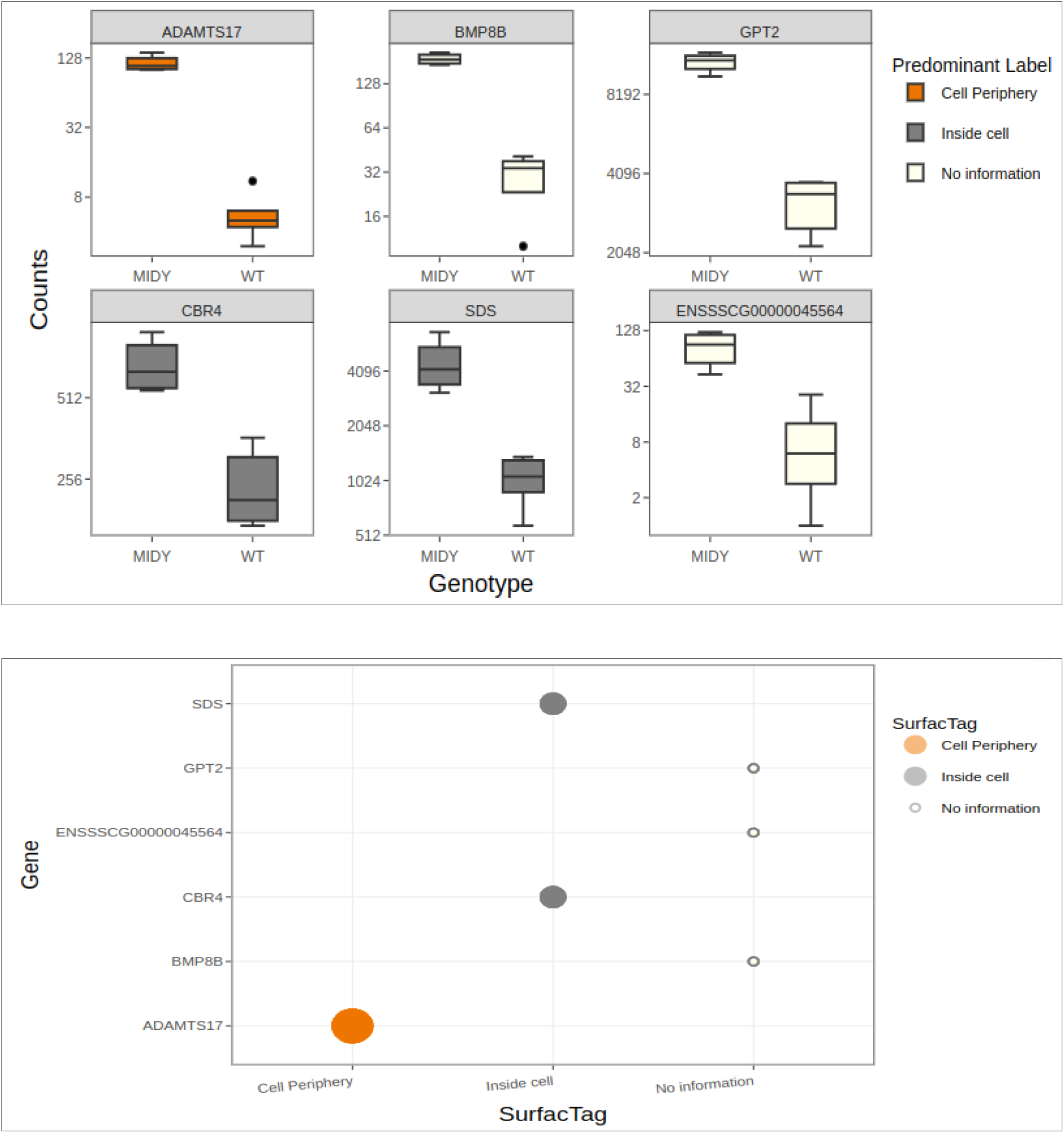
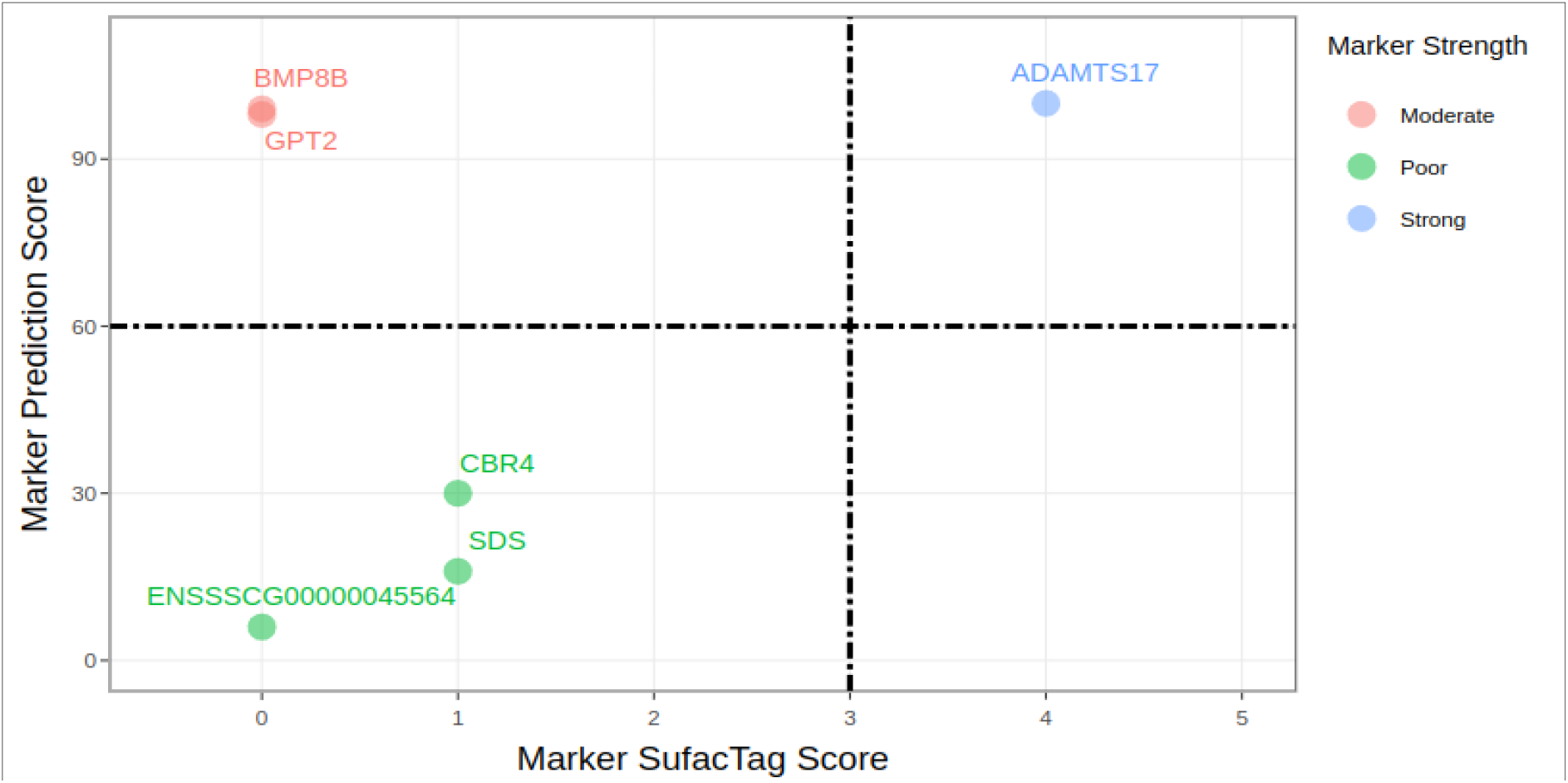
Visualization of Biomarker Analysis Results. **(A)** Boxplots displaying the differential expression of *SurfacOmics*-identified biomarkers between MIDY and WT groups. **(B).** Bubble plot illustrating cellular localization and labels from SurfacTag knowledgebase for identified biomarkers. **(C)**. *SurfacOmics* marker potential plot, highlighting ADAMTS17 as a candidate with substantial strength and elevated marker potential.

### *SurfacOmics* successfully identified biomarkers from Transcriptomics data

*SurfacOmics* provides a pertinent list of potential biomarkers for distinguishing the targeted conditions (genotype, i.e., learning variable). Crucial biomarkers identified by *SurfacOmics* are *ADAMTS17, BMP*8B, *CBR4, GPT2* and *SDS*. However, *ADAMTS17* had the highest Marker Potential score with decent cellular localization and a good prediction score **(Table 1)**.

For ease of visualization, *SurfacOmics* also comes with various plots such as box plots (MPS) and bubble plots (MSS, Marker potential). For instance, **Figure 3A** box plot depicts a comparable expression of *ADAMTS17* between WT and MIDY groups. Similarly, **Figure 3B** plot displays an additional layer of information in terms of cellular localization, identifying *ADAMTS17* on the cell surface, making it an easily assayable biomarker. Lastly, **Figure 3C** reflects the Marker Potential of each identified marker as a weighted sum of both MPS and MSS.

Interestingly, these results were found to be aligned with the original paper findings. Similarly on a proteomics level we have identified Q8WNW3, Q2QLE3, Q06AS6, P37111 and A0A286ZN52 **(Supplementary data)**. These concordant results focus on the importance of statistical stringency and easy-to-use *in-silico* tools for the identification of biomarkers. Here, we present *SurfacOmics*, that is designed to predict the biomarkers in six models i,e., *Macaca mulatta, Mus musculus, Sus scrofa, Danio rerio*, and *Drosophila melanogaster*. It combines the strength of solid statistical knowledge, equipped with manually curated knowledgebase in sync with biological parameters to achieve the set of potential biomarkers candidates.

## Conclusions

We present a user-friendly R-based GUI application *SurfacOmics*, developed to assist wet lab scientists and clinical researchers. This tool aims to enable prediction, prioritization, and ranking cell surface biomarkers from transcriptomics and proteomics datasets. It is equipped with advanced analytics and an accessible user interface, that enables quick data uploads and easy to interpret visualizations, simplifying the biomarker discovery process.

Biomarkers are identified using a penalized regression-based machine learning model and further annotated via SurfacTag knowledgebase. Additionally, results are presented in a downloadable tabular format with respective scores and additional information. The additional information highlights the biological significance of the prominent biomarkers and offers link-outs for antibody resources to simplify downstream analysis.

In conclusion, this tool allows users to predict and prioritize potential biomarker candidates that can significantly impact clinical research and enhance translational research. It has the ability to optimize the selection of genes or proteins for further investigation and intervention, thereby advancing our comprehension of cellular function and enhancing translational outcomes. Furthermore, we suggest wet lab-based studies and procedures using *in vitro* and *in vivo* models to confirm the safety and potency of the predicted biomarkers.

## Limitations

The current version of *SurfacOmics* is applicable for transcriptomics and proteomics datasets only. We aim to touch upon other omics domains and integrate it with significant clinical information.

## Data availability

Data will be made available on request.

## Acknowledgements & Funding

We are thankful to the Tata Consultancy Services (TCS) for providing an infrastructure support & funding.

## Notes

### Competing Interest Statement

The authors have declared no competing interest.

