## Supplemental Data for "*SurfacOmics:* an R shiny application integrating variable Feature Selection for gene biomarker discovery using Elastic-Net Regularization"

### SurfacOmics
### Supplementary Data

**SurfacTag Knowledgebase-SurfacTag**

Labels has been assigned to categorize and interpret genes and proteins based on their cellular component characteristics. They enable researchers to classify markers into localization groups, simplifying approach of cellular localization, henceforth accesibility and assaybility. These labels aid in hypothesis generation, experimental design, and the prioritization of research efforts. They are instrumental in advancing our understanding of cellular processes, facilitating drug discovery, and guiding studies aimed at uncovering the cellular mechanisms underlying health and disease.

Following are labels:
**1. Inside Cell:** The label "Inside cell" is assigned to genes and proteins whose primary location or function is situated within the cellular confines.

**2. Outside Cell:** The proteins or genes that are secreted or located in the extracellular environment outside the cell. These genes are often associated with intercellular communication, cell adhesion, or secretion of substances into the extracellular matrix.

**3. On Cell Surface:** Gene's products, such as proteins or receptors, are localized on the cell's outer membrane or surface. Genes with this label are frequently involved in cell signaling, interactions with neighboring cells, or environmental sensing.

**4. Outside on Cell Surface:** The label "Outside on cell surface" is applied to genes and proteins that occupy both extracellular spaces and the outer surface of the cell membrane. These molecules are involved in complex processes that bridge the intracellular and extracellular environments. The label is particularly useful in highlighting genes with versatile roles, including cellular communication, adhesion, and immune responses.

**5. Functional Dependent:** "Functional Dependent" is a label assigned to genes and proteins whose activity or function relies heavily on specific contextual cues or conditions. This label acknowledges that a gene's behaviour may vary in response to different biological contexts, such as cell type, developmental stage, or environmental factors.

**6. Not Clear:** The "Not Clear" label is a catch-all category often used when the precise function or localization of the gene is not well-defined or is subject to further research.

Utilizing the assigned labels and their respective frequencies in association with genes and proteins, a set of distinct use cases has been meticulously crafted to ascertain the predominant label accurately. The predominant label is termed as SurfacTag. Following are the use case:

**Supplementary Table 1. Use case for determining SurfaceTag redefined on top of Cellular Compartment labels**

| Cases | Protein | Label | Frequency | SurfaceTag |
|---|---|---|---|---|
| **Case 1** | Q28DL4 | Inside cell | 3 | Inside cell |
| **Case 2** | O75400 | Function Dependent | 1 | Function Dependent |
| | O75400 | Inside cell | 5 | Function Dependent |
| | O75400 | On cell surface | 1 | Function Dependent |
| **Case 3** | Q2G0U9 | On cell surface | 2 | On cell surface |
| | Q2G0U9 | Outside cell | 1 | On cell surface |
| | Q2G0U9 | Outside on cell surface | 1 | On cell surface |
| **Case 4** | Q2I0M5 | On cell surface | 1 | Cell Periphery |
| | Q2I0M5 | Outside cell | 1 | Cell Periphery |
| **Case 5** | Q2KHT4 | Inside cell | 1 | Function Dependent |
| | Q2KHT4 | On cell surface | 1 | Function Dependent |

**1: Single Label (Case 1):** If a gene has only one label associated with it (e.g., "Inside cell"), the SurfacTag for that gene is the same as the single label it possesses. In this case, the label is clear and unambiguous.

**2: Functional Dominance (Case 2):** When a gene is labeled as "Functional Dependent," regardless of the presence of other labels, the predominant label is set to "Functional." This simplifies the classification, emphasizing the gene's dependency on specific conditions or factors for its function.

**3:Highest Frequency (Case 3):** If a gene has multiple labels, and one of those labels has a higher frequency compared to others (e.g., "Inside cell" appears three times while other labels appear once), the label with the highest frequency becomes the SurfacTag.

4: E**qual Frequency - Cell Periphery (Case 4):** In cases where a gene has multiple labels with equal frequency, and those labels belong to the "Cell Periphery" category (e.g., "On cell surface" and "Outside on cell surface" both occur twice), the predominant label is set as "Cell Periphery." This reflects the gene's involvement in cell periphery region.

5: **Functional with Inside Cell (Case 5):** When a gene labelled as "Inside cell" shares equal frequency with labels from the "Cell Periphery" category (e.g., "Inside cell" and "On cell surface" both occur twice with equal frequency), the SurfacTag is set to "Functional." This reflects the gene's dual involvement in both intracellular and cell periphery activities.

**Marker SurfacTag Score (MSS)**

**Supplementary Table 2. MSS-based labels used to assign the cellular annotation.**
MSS is scoring matrix used as prioritizing measure to determine the cellular localization for individual gene and protein based on SurfacTag Labels.

| Cellular Annotation | MSS |
|---|---|
| No information | 0 |
| Inside cell | 1 |
| Outside cell | 2 |
| Function dependent | 3 |
| On cell surface | 4 |
| Outside on the cell surface | 4 |
| Cell Periphery | 4 |

**Results on Proteomic dataset**

***SurfacOmics* identified biomarkers from proteomic data**
*SurfacOmics* provides pertinent list of markers with potential for distinguishing the targeted conditions (Genotype). Markers identified by *SurfacOmics* are Q8WNW3, Q06AS6, A0A286ZN52, A0A286ZPR8 and R4HZ39.

**Table 3. Proteomic Biomarkers and corresponding scores identified by *SurfacOmics* application.**

| Marker Protein | Marker Prediction Score | Marker Surface Score | Marker Potential | Marker Strength |
|---|---|---|---|---|
| Q8WNW3 | 57 | 3 | 0.642 | Moderate |
| Q06AS6 | 100 | 0 | 0.6 | Moderate |
| A0A286ZN52 | 97 | 0 | 0.582 | Moderate |
| A0A286ZPR8 | 85 | 0 | 0.51 | Moderate |
| R4HZ39 | 73 | 0 | 0.438 | Moderate |
| A0A0K1TQQ7 | 59 | 0 | 0.354 | Poor |
| A0A286ZPD7 | 54 | 0 | 0.324 | Poor |
| Q06AA9 | 25 | 1 | 0.25 | Poor |
| Q06AB3 | 20 | 1 | 0.22 | Poor |
| Q06A99 | 20 | 0 | 0.12 | Poor |
| Q06AA0 | 20 | 0 | 0.12 | Poor |
| A0A286ZL65 | 17 | 0 | 0.102 | Poor |

For better visualization, SurfacOmics comes with various plots such as box-plots (MPS), bubble plot (MSS, Marker potential). For instance, **Figure 1A)** box plot depicts a good expression variation of identified biomarkers between WT and MIDY groups. **Figure 1B)** plot displays an additional layer of information in terms of cellular localization, identifying Q8WNW3 as Function Dependent. **Figure 1C)** reflects the Marker Strength/Potential of each identified marker as a weighted sum of both MPS and MSS.

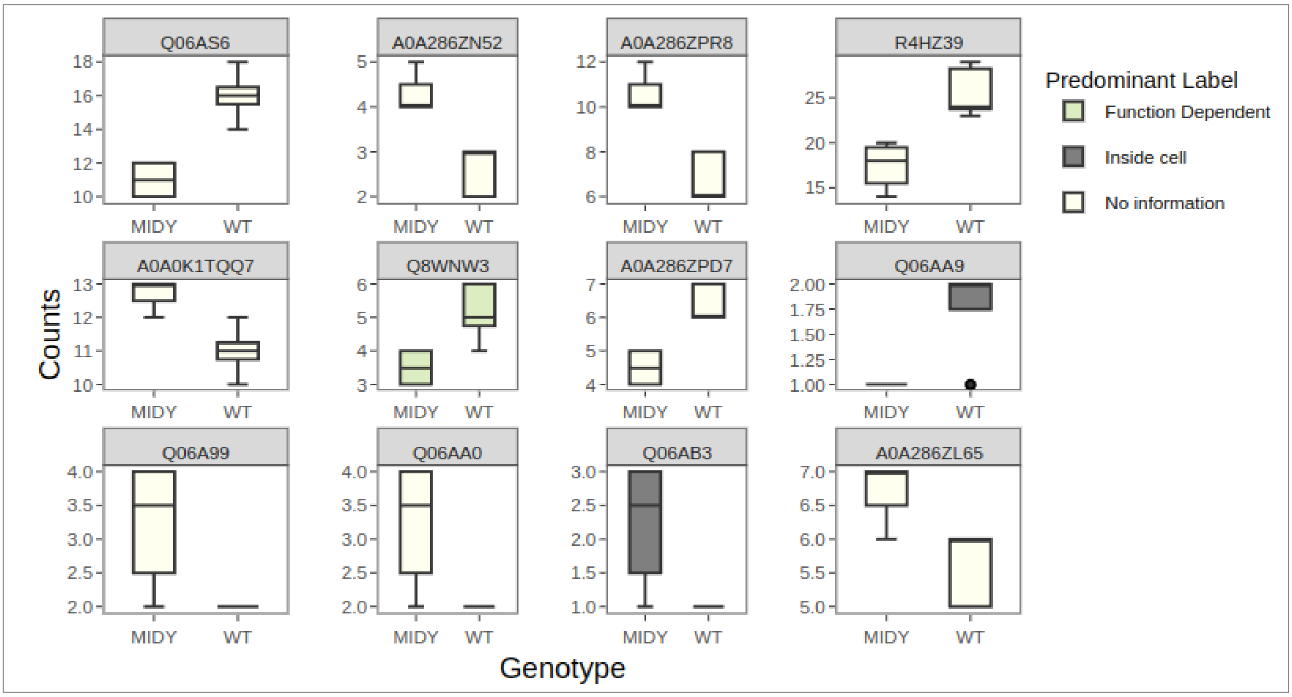


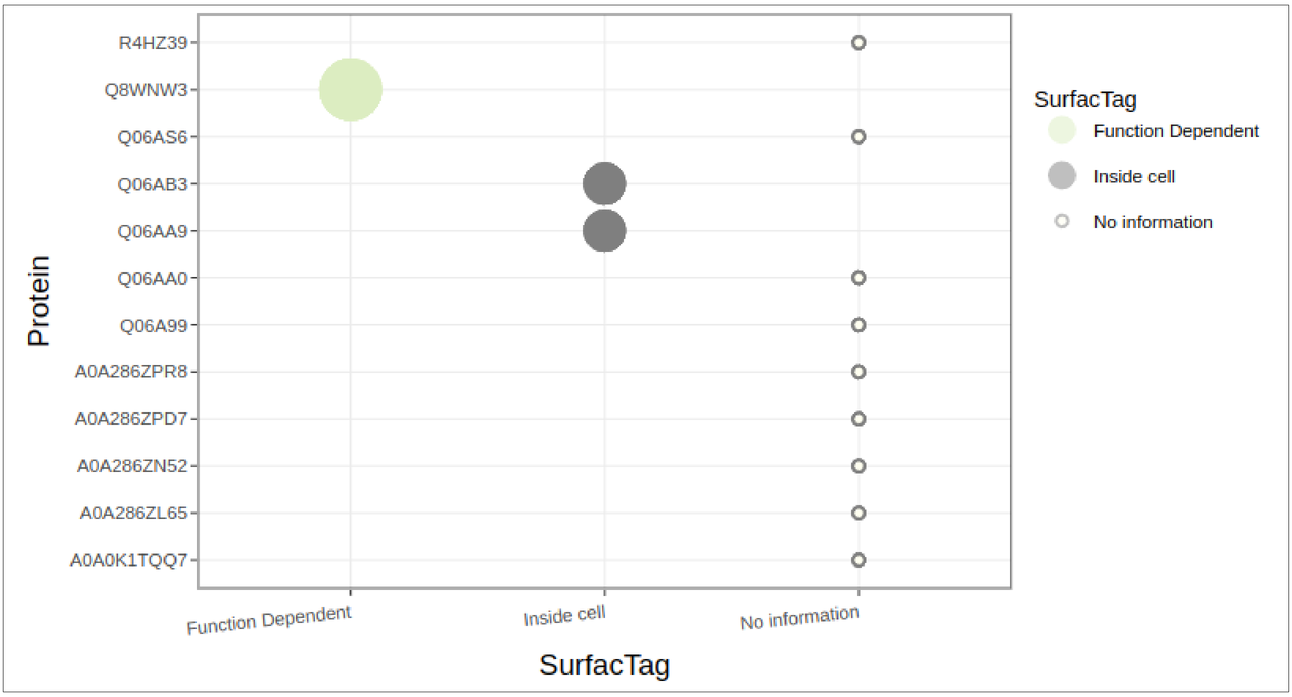

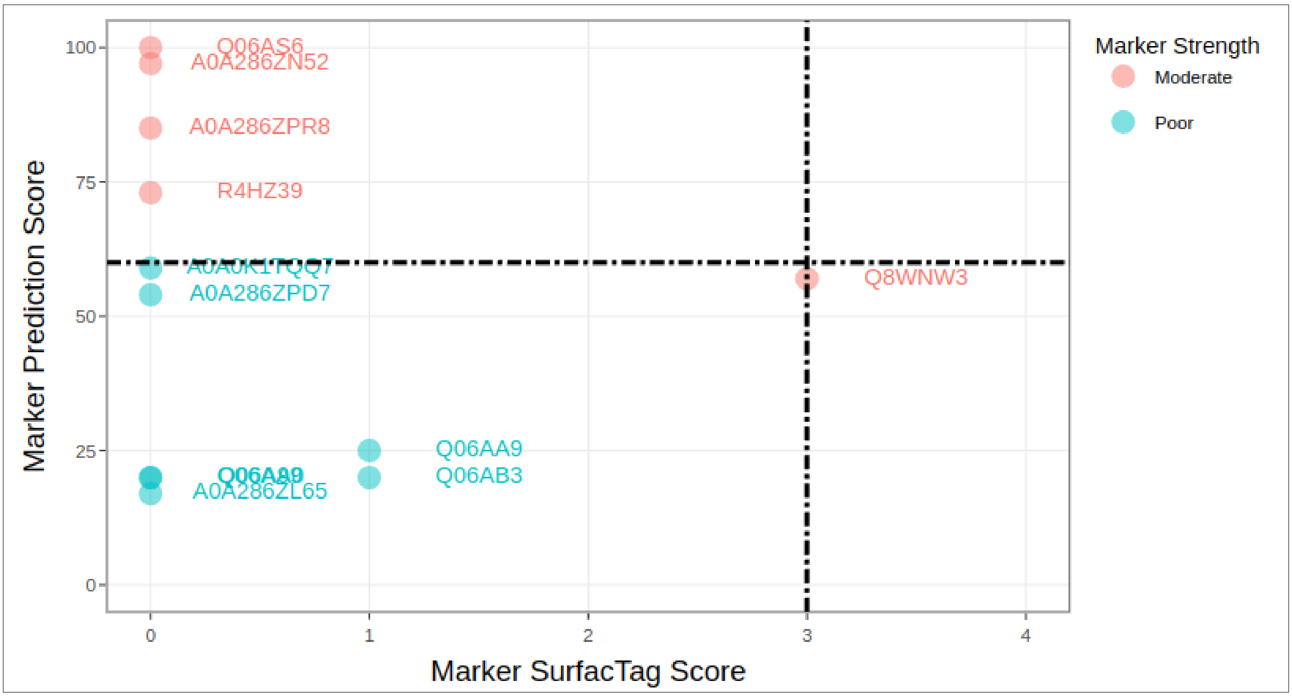


**Figure 1. (A)** Box plots showing the expression difference of SurfacOmics-identified biomarkers between MIDY and WT groups. **(B).** Bubble plot showing cellular localization and labels from knowledge-base for identified biomarkers. **(C).** *SurfacOmics* marker potential plot, showing Q8WNW3 as Function Dependent biomarkers.

To provide a comprehensive view, SurfacOmics provides annotated information encompassing molecular functions, biological processes and antibody resource linkouts associated with each gene and protein. Following

**Table 4.** summarizes comprehensive information of proteomics marker in tabular format.

| Protein | Predominant_ Label | Protein Names | Gene Names | Molecular Function | Biological Process | Antibody_Resource |
|---|---|---|---|---|---|---|
| A0A0K1TQ Q7 | No information | No information | No information | No information | No information | https://www.antibody resource.com/proteins earch/Protein/?search Term=A0A0K1TQQ7 |
| A0A286ZL65 | No information | No information | No information | No information | No information | https://www.antibody resource.com/proteins earch/Protein/?search Term=A0A286ZL65 |
| A0A286ZN5 2 | No information | No information | No information | No information | No information | https://www.antibody resource.com/proteins earch/Protein/?search Term=A0A286ZN52 |
| A0A286ZPD 7 | No information | No information | No information | No information | No information | https://www.antibody resource.com/proteins earch/Protein/?search Term=A0A286ZPD7 |
| A0A286ZPR 8 | No information | No information | No information | No information | No information | https://www.antibody resource.com/proteins earch/Protein/?search Term=A0A286ZPR8 |
| Q06A99 | No information | No information | No information | No information | No information | https://www.antibody resource.com/proteins earch/Protein/?search Term=Q06A99 |
| Q06AA0 | No information | No information | No information | No information | No information | https://www.antibody resource.com/proteins |

| | | | | | | |
|---|---|---|---|---|---|---|
| | | | | | | earch/Protein/?search Term=Q06AA0 |
| Q06AA9 | Inside cell | Ubiquitin-conjugating enzyme E2 D2 (EC 2.3.2.23) ((E3-independent) E2 ubiquitin-conjugating enzyme D2) (EC 2.3.2.24) (E2 ubiquitin-conjugating enzyme D2) (Ubiquitin carrier protein D2) (Ubiquitin-protein ligase D2) | UBE2D2 UBC4 UBCH4 UBCH5B | ATP binding ; ubiquitin conjugating enzyme activity ; ubiquitin protein ligase binding ; ubiquitin-protein transferase activity | protein K11-linked ubiquitination ; protein K48-linked ubiquitination ; ubiquitin-dependent protein catabolic process | https://www.antibodyresource.com/proteinsearch/Protein/?searchTerm=Q06AA9 |
| Q06AB3 | Inside cell | Ubiquitin carboxyl-terminal hydrolase isozyme L3 (UCH-L3) (EC 3.4.19.12) (Ubiquitin thioesterase L3) | UCHL3 | cysteine-type deubiquitinase activity | protein deubiquitination ; ubiquitin-dependent protein catabolic process | https://www.antibodyresource.com/proteinsearch/Protein/?searchTerm=Q06AB3 |
| Q06AS6 | No information | No information | No information | No information | No information | https://www.antibodyresource.com/proteinsearch/Protein/?searchTerm=Q06AS6 |
| Q8WNW3 | Function Dependent | Junction plakoglobin | Jup | alpha-catenin binding ; cadherin binding ; DNA-binding transcription factor binding ; protein phosphatase binding ; transcription coactivator activity | canonical Wnt signaling pathway ; cell-cell adhesion ; positive regulation of transcription by RNA polymerase II | https://www.antibodyresource.com/proteinsearch/Protein/?searchTerm=Q8WNW3 |
| R4HZ39 | No information | No information | No information | No information | No information | https://www.antibodyresource.com/proteinsearch/Protein/?searchTerm=R4HZ39 |
